# Two RhoGEF isoforms with distinct localisation act in concert to control asymmetric cell division

**DOI:** 10.1101/2022.11.06.515358

**Authors:** Emilie Montembault, Irène Deduyer, Marie-Charlotte Claverie, Lou Bouit, Nicolas Tourasse, Denis Dupuy, Derek McCusker, Anne Royou

**Author notes:** Contributed equally (Emilie Montembault, Irène Deduyer, Marie-Charlotte Claverie).

## Abstract

Cytokinesis is essential for the partitioning of cellular contents into daughter cells. It relies on the formation of an acto-myosin contractile ring, whose constriction induces the ingression of the cleavage furrow between the segregated chromatids. Rho1 GTPase and its RhoGEF (Pbl) are essential for this process as they drive the assembly and constriction of the contractile ring. However, how Rho1 is regulated to sustain efficient furrow ingression while maintaining correct furrow position remains poorly defined. Here, we show that during asymmetric division of Drosophila neuroblasts, Rho1 is controlled by two Pbl isoforms with distinct localisation. Spindle midzone- and furrow-enriched Pbl-A focuses Rho1 at the furrow to sustain efficient ingression, while Pbl-B pan-plasma membrane localization promotes the broadening of Rho1 activity and the subsequent enrichment of cortical myosin. This enlarged zone of Rho1 activity becomes essential to adjust furrow position during ingression, thereby preserving correct daughter cell size asymmetry. Our work highlights how the use of isoforms with distinct localisation patterns provides robustness to an essential process.

## Introduction

Cell cleavage, also called cytokinesis, is a fundamental process that occurs subsequent to sister chromatid segregation. In animal cells, cytokinesis relies on the assembly of a contractile ring composed of filamentous actin and bipolar filaments of non-muscle myosin II (referred to as myosin) anchored at the equatorial plasma membrane. Myosin activity generates the force necessary to drive the ingression of the cleavage furrow. The topological resolution of the two resulting cells is achieved with abscission of the membrane (D’Avino et al., 2015; Green et al., 2012). The activity of the small GTPase RhoA (Rho1 in Drosophila) is essential for this process as it promotes actin nucleation and myosin activation (Dean et al., 2005; Kishi et al., 1993; Pollard and O’Shaughnessy, 2019; Prokopenko et al., 1999). Rho1 is a small GTPase of the Ras super family that undergoes nucleotide-induced conformational changes as it toggles between inactive GDP- and active GTP-bound states. Rho1 localizes to the plasma membrane through prenylation at its carboxy-terminus and its lipid interaction is obligatory for function (Allal et al., 2000). The zone of active Rho1 at the plasma membrane determines the position of contractile ring assembly and subsequent cleavage furrow ingression (Bement et al., 2005; Wagner and Glotzer, 2016). Rho1 activation is catalyzed by a guanine nucleotide exchange factor (GEF) called Ect2 (Epithelial transforming 2) in mammals (Basant and Glotzer, 2018; Kim et al., 2005; Miki et al., 1993; Tatsumoto et al., 1999) and Pebble (Pbl) in drosophila (Hime and Saint, 1992; Lehner, 1992; Prokopenko et al., 1999). Hence, the mechanism controlling Ect2/Pbl localisation and activation is critical to determine the zone of Rho1 activation and thus the position of the resulting furrow. Ect2/Pbl contains BRCT (BRCA1 C-terminal) domains in its N-terminus and a catalytic DH (Dbl homology) domain and juxtaposed membrane-associated PH (Plekstrin homology) domain in its C-terminus. The canonical mechanism by which Ect2/Pbl concentrates at the equatorial zone to activate Rho1 is through its interaction with microtubule-associated mgcRacGAP/Cyk4 (RacGAP50C or tumbleweed in Drosophila) (Goldstein et al., 2005; Somers and Saint, 2003; Yuce et al., 2005). RacGAP50C forms part of the conserved centralspindlin complex with the kinesin 6 MKLP1/Zen4 (Pavarotti in Drosophila). This complex promotes the bundling of anti-parallel microtubules at their plus-end to form the midzone of the central spindle that assembles between the two segregated pools of chromatids (Gomez-Cavazos et al., 2020; Hime and Saint, 1992; Somers and Saint, 2003; Tatsumoto et al., 1999; Yuce et al., 2005; Zavortink et al., 2005). The phosphorylation of RacGAP50C by Polo-like Kinase 1 (Polo in Drosophila) provides docking sites for Ect2/Pbl’s BRCT domains, thereby relieving Ect2/Pbl auto-inhibition and promoting its accumulation at the spindle midzone and equatorial membrane (Adams et al., 1998; Burkard et al., 2007; Gomez-Cavazos et al., 2020; Schneid et al., 2021; Wolfe et al., 2009).

While the molecular pathway that specifies the zone of Rho1 activation to initiate contractile ring assembly is well defined, less is known about the mechanisms that maintain the contractile ring position while sustaining efficient constriction. The study of these mechanisms is hindered by the fact that severe perturbation of either Rho1 signalling or the kinetics of contractile ring components during division results in failure to initiate furrowing or furrow regression. However, the mechanism that controls Rho1 activity to sustain furrowing at the right position is critical to preserve the size of the resulting daughter cells. This is particularly important during asymmetric stem cell division where cell size determines cell fate, as observed in the Drosophila pIIb sensory organ precursor lineage and the *C. elegans* Q neuroblast lineage, where the smaller daughter cells undergoes apoptosis (Fichelson and Gho, 2003; Ou et al., 2010).

The Drosophila larval neuroblast is a powerful model for studying the mechanisms that spatio-temporally control the asymmetric position of the furrow. The division of this large stem cell produces two cells that differ in size and fate: the larger cell retains the neuroblast “stemness” and continues dividing frequently, while the smaller ganglion mother cell (GMC) undergoes one round of division before differentiating into neuronal lineages (Neumuller and Knoblich, 2009). Previous studies have shown that the asymmetric position of the furrow is specified by two parallel pathways, a spindle-independent pathway that requires the polarity complex Pins (composed of Partner of Inscuteable (Pins); Gαi; Discs Large) to bias Myosin activity towards the basal cortex and a microtubules-dependent pathway that regulates RacGAP50C localisation to the equatorial cortex (Cabernard et al., 2010; Cai et al., 2003; Connell et al., 2011; Giansanti et al., 2001; Pham et al., 2019; Roth et al., 2015; Roubinet et al., 2017; Thomas et al., 2021; Yu et al., 2003).

In addition to these pathways, our team recently identified novel Pbl- dependent myosin dynamics during neuroblast division. At mid-ring closure, a pool of myosin undergoes outward flow (efflux) from the contractile ring, invading the entire cortex (hereafter referred to as the “polar” cortex or the “poles” as opposed to the “equatorial” cortex where the furrow forms) until the end of furrow ingression (Montembault et al., 2017). This pool of active myosin at the poles becomes critical if chromatids trail at the cleavage site during furrow ingression. Under these conditions, cell elongation ensues from the reorganization of polar myosin into broad lateral rings that partially constrict. This “adaptive” cell elongation facilitates the clearance of trailing chromatids from the cleavage site, thereby preventing chromatid arms being trapped by the contractile ring. Attenuation of Pbl function using the *pbl^MS^* hypomorphic allele in combination with a *pbl^5^* null mutation, impairs myosin efflux from the furrow to the poles during division. Consequently, cells cannot undergo adaptive elongation in the presence of trailing chromatids at the site of cleavage, with dire consequences for tissue homeostasis (Kotadia et al., 2012; Montembault et al., 2017). While these Pbl-dependent changes in the mechanical properties of the polar cortex are important for the clearance of trailing chromatids, little is known about their role in maintaining the fidelity of asymmetric neuroblast division under normal conditions.

In this study, we investigate the consequences of perturbed myosin polar cortex enrichment due to reduced Pbl activity during asymmetric cell division by live imaging of neuroblasts.

## Results

To determine the importance of myosin and, hence, Rho1 activity at the poles during asymmetric cell division, we examined myosin dynamics during drosophila larval neuroblast cytokinesis in wild type (WT) and in the *pbl^MS^*homozygote mutant. Neuroblasts expressing the non-muscle myosin II regulatory light chain (encoded by *spaghetti-squash*) fused to green fluorescent protein (referred to as myosin) were monitored by spinning-disk microscopy (Royou et al., 2002). In wild type neuroblasts, myosin adopted a uniform cortical localisation at anaphase onset (initiation of sister chromatid separation)(Figure 1A, video 1). Subsequently, myosin depleted the apical followed by the basal pole and enriched at the baso-lateral cortex to initiate furrowing (Figure 1A and S1) (Cabernard et al., 2010; Connell et al., 2011; Montembault et al., 2017; Roubinet et al., 2017). At mid-furrow ingression, a pool of myosin underwent outward flow from the ring and enriched the entire cortex of both nascent daughter cells (Figure 1A) (Montembault et al., 2017). Myosin concentration at the poles was maximal at 80% of furrow ingression (Figure 1A-E). Then, myosin was entirely dissociated from the cortex by the end of nuclear envelope reformation, as illustrated by the depletion of myosin from the nucleus (Figure 1A)(Montembault et al., 2017). In contrast, no myosin signal was detected at the polar cortex at any time points during cytokinesis in the *pbl^MS^* homozygote mutant, similar to the *pbl^5^/pbl^MS^* transheterozygote mutant (Figure 1A-E, video 1)(Montembault et al., 2017). After apical and basal depletion, Myosin rapidly compressed into a tight ring at the baso- lateral position, where it remained focused throughout closure (Figure 1A-E, Figure S1).

**Figure 1.**
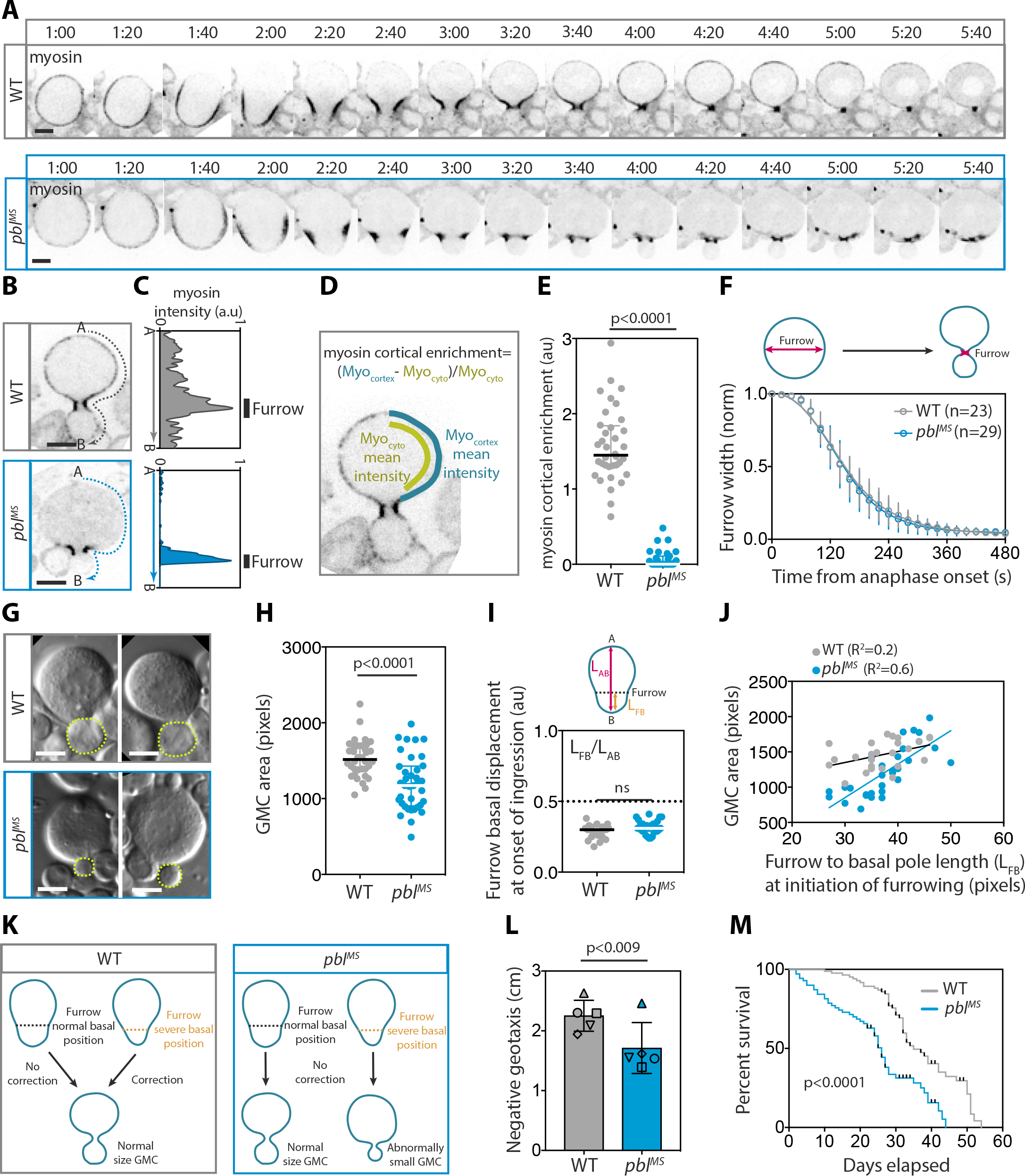
Defects in cortical myosin dynamics and daughter cell size asymmetry during cytokinesis in *pbl^MS^* mutant neuroblasts. (**A**) Time-lapse images of wild type (WT) and *pbl^MS^* mutant third instar larvae neuroblasts expressing the regulatory light chain of the non-muscle myosin II fused with GFP (Sqh::GFP, referred to as myosin). Time, min:sec. Time starts at anaphase onset. (**B**) Selected sagittal images of cells of the indicated genotype at a similar timepoint during cytokinesis and (**C**) the corresponding linescan of myosin average intensity at the cortex from apical (A) to basal (B) poles. Time is from anaphase onset. The average intensity of cytoplasmic myosin has been subtracted and the data normalized to the maximum intensity. Bars indicate median±interquartile range. (**D**) Scheme showing the method used to quantify the enrichment of myosin at the polar cortex and (**E**) the corresponding scatter dot plot. (**F**) Graph showing the furrow width over time for the indicated genotypes. n, number of cells. The yellow area corresponds to the average duration of myosin global cortical enrichment in WT cells. Symbols and bars represent mean±SD. (**G**) Differential interference contrast images of two distinct neuroblasts for each genotype at the end of cytokinesis. The dashed yellow lines delineate the GMC area. (**H**) Scatter dot plot showing the GMC area (number of pixels) for the indicated genotypes. (**I**) Scatter dot plot showing the basal displacement of the furrow at onset of ingression for the indicated genotypes calculated as illustrated in the scheme above the graph. For all scatter dot plots, bars correspond to median ± interquartile range. A Mann-Whitney test was used to calculate P values. (**J**) Graph showing the correlation between the basal position of the furrow at the onset of ingression and the size of the resulting GMC at the end of ingression for the indicated genotype. Pearson r correlation coefficient was used to calculate R^2^. A significant positive correlation is found in *pbl^MS^* mutants (P<0.0001) but not in WT. (**K**) Scheme illustrating the model for the abnormally small GMC phenotype resulting from *pbl^MS^* mutant neuroblast divisions. In WT and *pbl^MS^* mutant cells, the furrow is positioned towards the basal pole at the onset of ingression. However, some furrows are positioned more basally than others. In WT cells, this severe basal position is corrected during ring constriction to preserve the size of the resulting GMC at the end of ingression. In contrast, no correction occurs in the *pbl^MS^* mutant, which means that a furrow with a severe basal position at the start of ingression will produce an abnormally small GMC. (**L**) Graph showing the negative geotaxis for the indicated genotypes. Histogram and bars represent mean±SD of 5 independent experiments. Symbols represent paired experiments. A one-tailed paired student t-test was used to calculate P value. (**M**) Kaplan-Meier survival curves of WT and *pbl^MS^*adults. The data represent five vials of 10 adults of equal gender each for the indicated genotype. *pbl^MS^* mutant have a significantly reduced longevity compared to wild-type strain (P<0.0001, Mantel-Cox test). Scale bars, 5µm for all images.

Next, we determined if the absence of active myosin at the polar cortex affected the rate of furrow ingression. To do so, we measured furrow width over time from anaphase onset until the end of invagination (Figure 1F). No difference in the rate of furrow ingression was observed between wild type and the *pbl^MS^* mutant, suggesting that the outward flow of myosin from the contractile ring and transient activity at the global cortex during ring constriction was not important for the efficient invagination of the furrow.

Next we examined if the position of the furrow was maintained during ring constriction in the *pbl^MS^* mutant. To do so we first measured the area of the resulting GMC at the end of ring closure. Remarkably, a significant proportion of *pbl^MS^* mutant cell divisions led to the production of abnormally small GMCs, suggesting a defect in furrow position during ingression (Figure 1G and H). To determine if the abnormally small GMC size was due to furrow mis-positioning at an early stage during cleavage, we measured its position at the onset of ingression. We found that, as previously reported, the furrow was positioned towards the basal pole in WT cells (Figure 1I) (Cabernard et al., 2010; Connell et al., 2011; Roth et al., 2015; Roubinet et al., 2017). No difference in initial furrow position was observed in *pbl^MS^* mutants compared to WT, indicating that the small GMC phenotype in the *pbl^MS^* mutant was not due to early mis-positioning of the furrow (Figure 1I). We noticed some variation in the initial basal position of the furrow. Some furrows were more basally positioned than others in both WT and *pbl^MS^* mutant cells. Since not all *pbl^MS^* divisions produce abnormally small GMCs, we reasoned that the initial basal position of the furrow determined the size of the resulting GMC in *pbl^MS^* mutants. To test this hypothesis, we plotted the position of the furrow at ingression onset (as the distance from the furrow to the basal pole) with the area of the resulting GMC. Remarkably, in contrast to WT cells, a strong correlation was found between the initial position of the furrow and the size of the resulting GMC in *pbl^MS^*mutants (Figure 1J). These results suggest a model in which, in WT cells, the variation in the initial position of the furrow is sensed so that severe basal positions are corrected during ring constriction, thereby preserving the size of the resulting GMC. This correction mechanism does not exist in the *pbl^MS^*mutant, hence furrows that are initially in an extreme basal position produce abnormally small GMCs (Figure 1K).

Changes in cell size can affect cell fate by inducing cell death (Chartier et al., 2021; Chen et al., 2016; Ou et al., 2010). Thus we reasoned that the abnormally small GMC observed in the *pbl^MS^* mutant might be eliminated during development, resulting in cognitive impairment. To test this hypothesis, we assessed locomotion behaviour using the Rapid Iterative Negative Geotaxis (RING) assay (Figure 1L)(Gargano et al., 2005). We found a significant reduction in negative geotaxis for the *pbl^MS^* mutant adults compared to WT, suggesting a locomotion decline in this mutant. Next, we assessed how the *pbl^MS^* mutation impacts adult longevity. *Pbl^MS^* is a viable mutation that induces male sterility due to cytokinesis failure in spermatocytes (Giansanti et al., 2004). Comparison of survival curves between wild type and the *pbl^MS^* mutant revealed a significant shortening of *pbl^MS^* adult lifespan (Figure 1M). Collectively these results suggest that the physiological consequences of abnormally small GMC production are their elimination during development with dire consequence for the organism.

The *pbl^MS^* allele was identified through an EMS screen for male sterility due to defects in spermatocyte division (Giansanti et al., 2004). However, the mutation responsible for the *pbl^MS^* phenotype has not yet been characterized. To begin to understand how Pbl regulates asymmetric cytokinesis, we sequenced all the exons of the *pbl* gene (CG8114) from *pbl^MS^* male adult genomic DNA extracts and compared these with the *pbl* sequence of the wild type strain we had in the lab. The Flybase database annotation reports that the *pbl* gene is encoded by 15 exons spread over 15 kb on chromosome 3L (cytologic position 66A18-66A9 and genomic position 7896109-7911954). We identified a cytosine to thymidine substitution in exon 8 at position 7901087 nucleotides on chromosome 3R in *pbl^MS^* DNA extracts from three independent experiments. This substitution changes the glutamine codon CAG into the stop codon TAG (Figure 2A).

**Figure 2.**
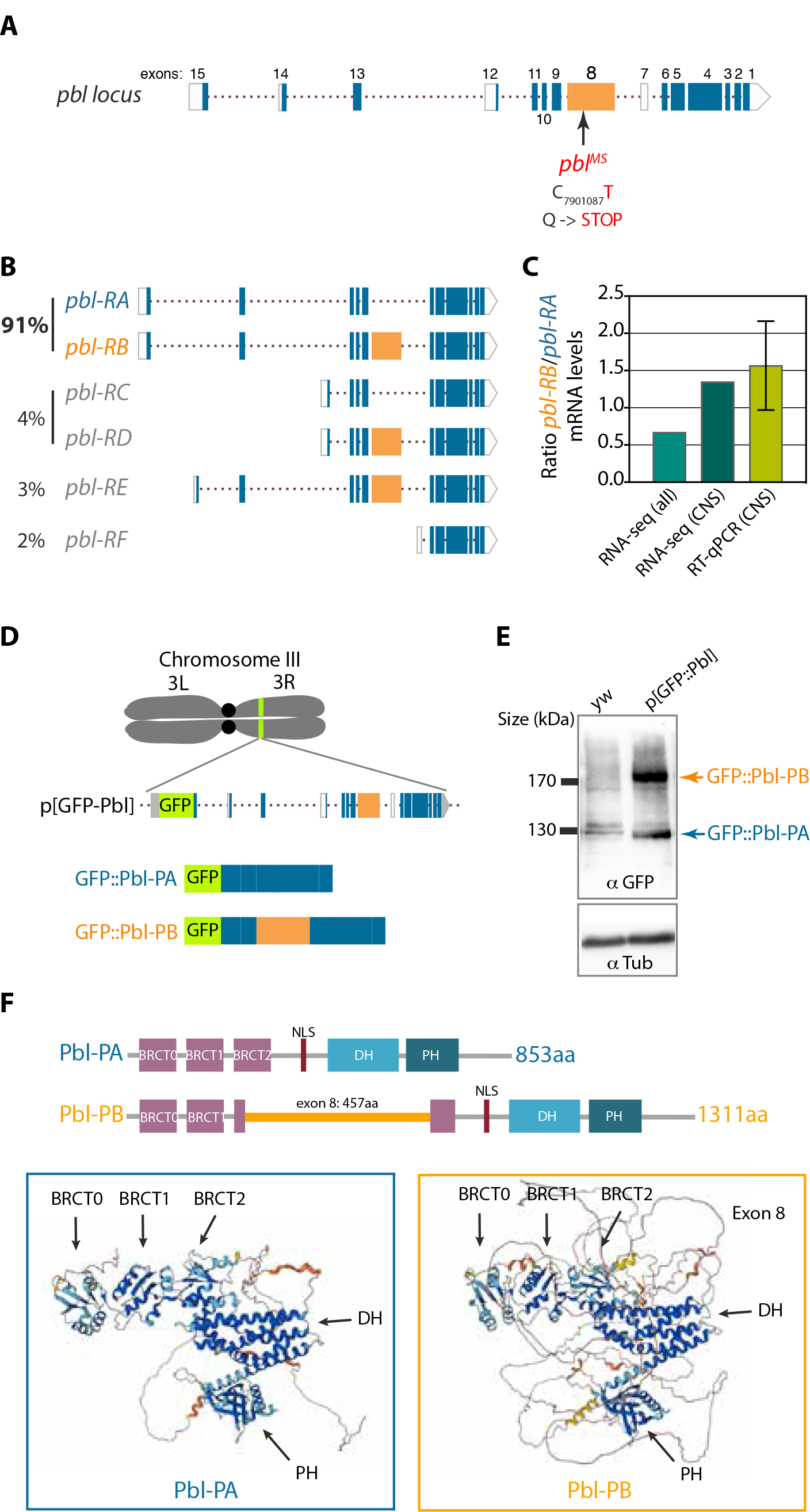
*pbl^MS^* is a non-sense substitution in an alternatively spliced exon specific to the Pbl-B isoform, which is expressed at equal levels as Pbl-A in the central nervous system. (A) Diagram of the *pbl* locus. Filled boxes represent coding regions (exon 8 is highlighted in orange) and empty boxes correspond to potential untranslated sequence. The arrow and number indicate the base substitution (C to T) in exon 8 and the position on the chromosome. This non-sense mutation generates a glutamine (Q) to stop codon in the alternatively spliced exon 8. (**B**) Diagrams of the six predicted *pbl* isoforms mRNAs from alternative splicing. The numbers represent the estimated frequencies of the pbl isoforms based on exon junction frequencies recovered from a compendium of RNA-seq experiments. The isoforms pbl-RA and pbl-RB account for 91% of expressed messengers at that locus. (**C**) Graph showing the Pbl-RB/Pbl-RA mRNA level ratio in the central nervous system using exon junction analysis from 1959 RNAseq dataset (95286 reads) (RNAseq all) and 13 central nervous system dataset (272 reads) (RNAseq CNS) as well as semi-quantitative RT-qPCR from CNS mRNA extracts. (**D**) Scheme of the p[GFP-Pbl] genomic construct inserted on the right arm of chromosome 3 used for the western blot shown in (E). (**E**) Western blot probed with anti-GFP antibodies to detect GFP::Pbl-A and GFP::B from the indicated genotypes (top panel). Detection of α-tubulin was used as a loading control (bottom panel). (**F**) Diagrams and AlphaFold structure predictions of Pbl-PA and Pbl-PB. Pbl-PA possesses three BRCT (BRCA1 Carboxyl terminal) domains in its N-terminus. The additional 457 amino acids of exon 8 are positioned within the third presumptive BRCT domain in Pbl-B. However, AlphaFold model predicts that the exon 8 amino acids form an unstructured loop that does not affect the folding of the third BRCT. Color code of AlphaFold model confidence: dark blue, very high, light blue, high, yellow, low, orange, very low.

The Flybase website reports that exon 8 is alternatively spliced and is present in two of the six predictive *pbl* protein isoforms, Pbl-B and D isoforms (uniprot: Q8IQ97)(Figure 2B). Hence, *pbl^MS^* mutant flies do not express functional Pbl-B and D. To determine the overall level of the six predicted *pbl* isoform mRNAs we retrieved the raw read sequences from 1959 RNA-seq experiments publicly available in the NCBI SRA database (https://www.ncbi.nlm.nih.gov/sra) and mapped them to the Drosophila genome using the software HISAT2 in order to quantify relative usage of all exon junctions of the *pbl* mRNA gene (Kim et al., 2015). The identification of exon junction usage was performed according to the methods described previously (Tourasse et al., 2017). This large-scale quantitative analysis revealed that the major Pbl mRNA isoforms detected overall and specifically in the central nervous system are pbl-RA and RB (Figure 2B). In addition, they are present at near similar amount in the central nervous system (Figure 2C). Next, we determined the relative mRNA levels of the two major *pbl* isoforms in the larval central nervous system. To do so we performed a semi-quantitative RT-PCR on mRNA samples extracted from third instar larval brains using primers that amplify Pbl-RA and RB. The ratio of Pbl-RB to Pbl-RA PCR product was calculated (Figure 2C). Our results revealed that both splicing variants A and B are expressed at near equal amount, consistent with our results from the Pbl exon junction usage analysis. In agreement, northern blots with Drosophila embryonic and adult mRNA extracts using *pbl* cDNA as a probe detected two *pbl* major transcripts of similar levels, a lower transcript corresponding to *pbl-RA* and a higher transcript consistent with the size of *pbl-RB* (Prokopenko et al., 2000).

Since mRNA levels do not necessarily correlate with the amount of protein they encode (Becker et al., 2018), we determined the amount of Pbl-A and B proteins in the larval brain. We were not able to detect both endogenous Pbl-A and B proteins from the same larval brain extracts by western blot using several batches of anti-Pbl antibodies. Therefore, to estimate the level of Pbl-A and B protein produced from alternative splicing in the larval brain we used transgenic flies containing Pbl genomic sequence including the GFP sequence inserted 5’ of the start codon and under the control of the endogenous *pbl* promoter (Figure 2D). Levels of GFP::Pbl-A and -B proteins were assessed by Western blot using anti-GFP antibodies on third instar larvae brain extracts (Figure 2E). Two bands corresponding to the expected size for GFP::Pbl-A and GFP::Pbl-B were detected only in the transgenic strain extract indicating that the two GFP::Pbl proteins are produced at similar levels from alternative splicing. It is thus likely that the two endogenous Pbl isoforms are expressed at equal levels in this tissue. Collectively, these results indicate that Pbl-A and -B are the two major Pbl splicing variants expressed overall. In addition they are present at similar levels in the central nervous system. In addition, since *pbl^MS^* cells do not express Pbl-B, it suggests that Pbl-B but not Pbl-A drives myosin polar cortex enrichment during cytokinesis and that Pbl-B is important to maintain the correct size of daughter cells during asymmetric cell division.

Next, we examined the differences in secondary structure predicted for Pbl-A and Pbl-B. Analysis of Pbl-A structure using the DeepMind software Alphafold predict a third BRCT domain (named BRCT0) in addition to the conserved N-terminal BRCT1 and 2 (Figure 2F) (https://alphafold.ebi.ac.uk/entry/Q9U7D8) (Senior et al., 2020). This is consistent with a recent phylogeny study on the functional evolution of BRCT domains, which identified a third BRCT in Ect2 N-terminus (Sheng et al., 2011). The structural analysis of the N-terminus of Ect2 confirmed the presence of three apposed BRCT domains (Zou et al., 2014). While there is evidence that two BRCT domains are involved in Pbl and human Ect2 interaction with RacGAP50C/MgcRacGAP, less is known about the role of the third BRCT in this interaction (Gomez-Cavazos et al., 2020; Prokopenko et al., 1999; Schneid et al., 2021; Wolfe et al., 2009; Zou et al., 2014). While no homology to other protein domains was identified in the 457 amino-acid sequence encoded by exon 8, we found that this sequence is inserted in the middle of the 76 amino acid region that is predicted to form the BRCT2 domain in Pbl-B (Figure 2F). Alphafold structural modelling of Pbl-B predicted that the sequence encoded within exon 8 is unstructured but does not seem to hinder the folding of the BRCT2 (https://alphafold.ebi.ac.uk/entry/Q8IQ97) (Figure 2F) (Senior et al., 2020).

The absence of cortical myosin enrichment observed in *pbl^MS^*cells lacking a functional Pbl-B suggests that Pbl-B spatially regulates Rho1 activity at the cortex differently than Pbl-A. Thus, we investigated if Pbl-B and Pbl-A exhibit distinct sub- cellular localisation during neuroblast cytokinesis. To do so we produced transgenic flies strains expressing GFP::Pbl-A or B under the endogenous *pbl* promoter (see material and method). We monitored GFP::Pbl-A or GFP::Pbl-B signals during cytokinesis in live neuroblasts and compared their localisation with Venus::RacGAP50C, the centralspindlin component known to interact with Pbl-A (Zavortink et al., 2005)(Figure 3, video 2). Recent studies have reported that, in addition to its localisation to the midzone, the centralspindlin component and RacGAP50C’s partner Pavarotti accumulates at the site of furrowing via its association with a sub-population of microtubules called peripheral astral microtubules (Thomas et al., 2021). In agreement with Pavarotti localisation, we detected Venus::RacGAP50C at the midzone and at the furrow site throughout ingression. Venus::RacGAP50C strongly labelled the midbody (the remnant of the contractile ring and midzone components) upon completion of ring closure. Previous work reported that Pbl-A concentrates at the furrow of dividing embryonic cells but is not detected at the midzone of the central spindle (Prokopenko et al., 1999). In contrast to embryonic cells, in dividing neuroblasts, GFP::Pbl-A was observed at the midzone from mid-anaphase until the completion of furrow ingression, concomitantly to its localisation at the furrow and polar membrane (Figure 3, video 2). Intriguingly, it accumulated more prominently at the neuroblast than the GMC membrane. Upon completion of furrow ingression, GFP::Pbl-A labelled the midbody and the plasma membrane of the nascent cells. GFP::Pbl-A was detected in the nucleus after completion of cleavage furrow ingression as reported previously, and was both in the nucleus and plasma membrane in interphase cells (data not shown)(Montembault et al., 2017). While GFP::Pbl-B shared a similar localisation at the polar membrane as GFP::Pbl-A during furrow ingression, striking differences in its dynamics were also observed. First, GFP-Pbl-B was neither detected at the midzone nor at the midbody. Second, it accumulated in the nucleus before completion of cleavage furrow ingression, prior to Pbl-A nuclear localisation, and was strictly nuclear at the end of cytokinesis and in interphase. Third, it was not enriched at the furrow (Figure 3, video 2). Importantly, GFP::Pbl-B dynamics were similar in the presence or absence of endogenous Pbl (Figure 3 and data not shown). This latter result indicates that the absence of Pbl-B at the midzone is not due to the titration of RacGAP50C by endogenous Pbl-A. Collectively, these results indicate that the presence of exon 8 promotes premature Pbl-B nuclear localisation and either inhibits its midzone localisation or favours its broad membrane localisation at the expense of the midzone.

**Figure 3.**
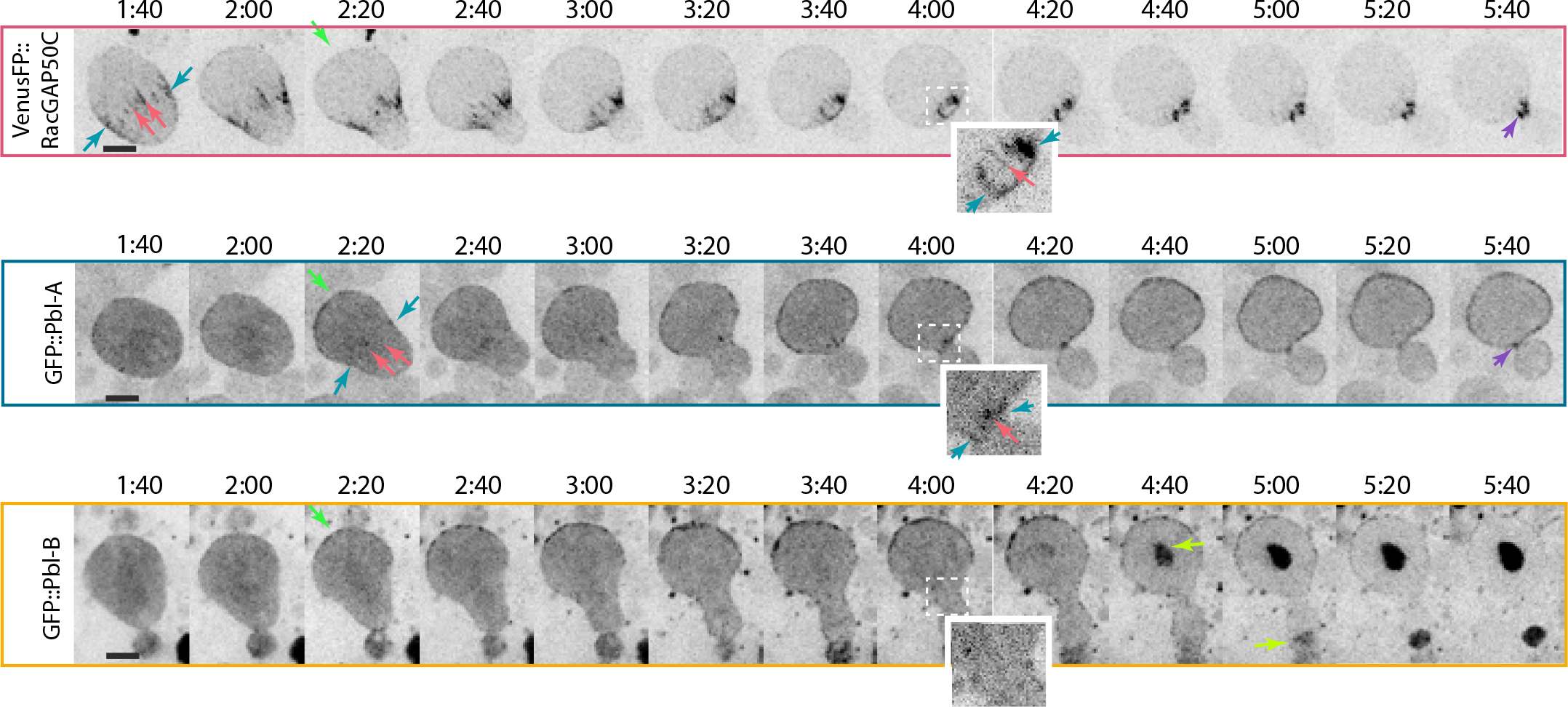
GFP::Pbl-A and GFP::Pbl-B have distinct localisation patterns during cytokinesis. Time-lapse images of neuroblasts expressing the indicated fluorescent proteins from anaphase B to late telophase. The green, blue, pink, yellow and purple arrows indicate the detection of the proteins at the plasma membrane, furrow, midzone, nucleus and midbody respectively. Insets are magnified images of the midzone and furrow region (indicated by the dashed square in the image). VenusFP::RacGAP50C and GFP::Pbl-A are detected at the midzone of the central spindle, at the furrow and polar membrane. In contrast, GFP::Pbl-B is only detected at the polar membrane but not at the midzone and furrow. It relocalizes to the nucleus prior to GFP::Pbl-A and is strictly nuclear at the end of cytokinesis. Note that both Pbl-A and B label more prominently the neuroblast than the GMC membrane. Time, min:sec, corresponds to the time from anaphase onset. Scale bar, 5µm.

The striking differences in Pbl-A and B localisation combined with the observation that myosin dynamics and cell size asymmetry during neuroblast divisions are affected in *pbl^MS^* mutants that lack a functional Pbl-B suggest that the two isoforms act differently to control Rho1 activity during cytokinesis. To examine the role of Pbl-A and B isoforms individually during neuroblast division, we adopted a genomic rescue approach. We took advantage of an existing pAcman plasmid containing a 17kb sequence including the *pbl* genomic sequence and the corresponding transgenic flies containing the *pbl* full-length construct (referred hereafter as Pbl-A+B) (Murray et al., 2012a). We produced two modified versions of this pAcman plasmid and the corresponding transgenic flies where the constructs were integrated at the same locus as the full-length to minimize variation in transgene expression. In the first version, we introduced a non-sense substitution in exon 8 mimicking the *pbl^MS^* mutation and thus preventing the production of a functional Pbl-B (referred hereafter as Pbl-A). In the second version, the intron between exon 8 and 9 was deleted to prevent alternative splicing of exon 8 and thus promote the expression of Pbl-B but not Pbl-A (referred hereafter as Pbl-B)(Figure 4A). The three constructs were first tested for their ability to rescue the lethality of the null *pbl* mutant transheterozygote *pbl^3^/pbl^2^* (Jurgens et al., 1984). All three constructs rescued the embryonic lethality of null *pbl* mutants. However, while Pbl-A+B and Pbl-A produced viable adult flies at the expected frequency according to the principles of Mendelian inheritance, the frequency of *pbl^3^/pbl^2^* hatching adults expressing only Pbl-B was dramatically reduced (Figure 4B). These results indicate that Pbl-A and B have redundant functions during development but Pbl-A either plays additional roles during development to adulthood or is required at a lower threshold than Pbl-B during development. Next we evaluated the efficiency of cytokinesis in the larval central nervous system by quantifying the frequency of polyploid mitotic cells on squashed brains that arise due to failed cytokinesis (Figure S2A and B). Pbl-A+B as well as Pbl-A larval brains exhibited negligible polyploid cells. Pbl-B larval brains displayed a mild increase in the frequency of polyploid cells per brain that was too mild, however, to account for the severe diminution of survival to adulthood.

**Figure 4.**
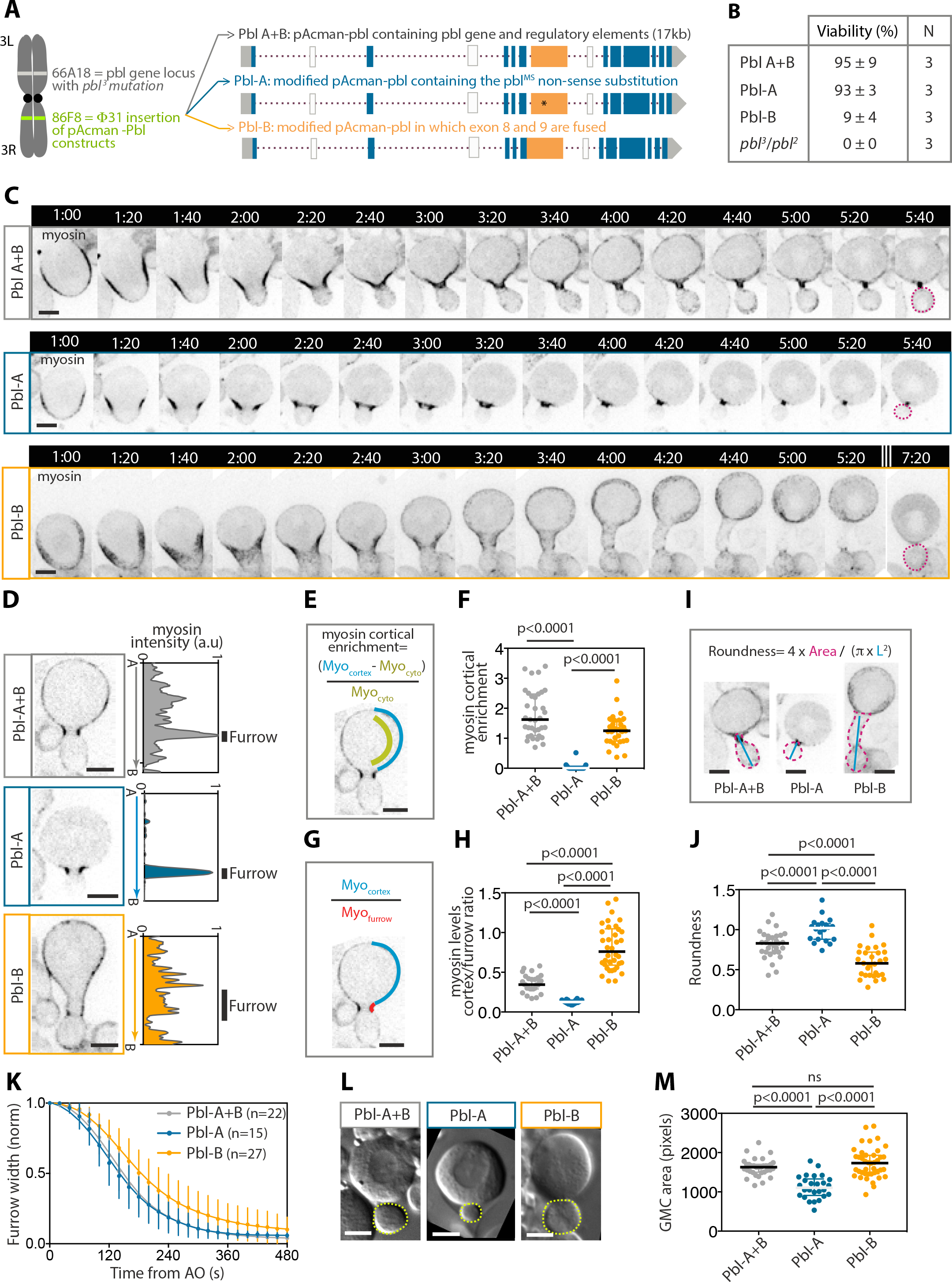
Pbl-A and B are both required for robust asymmetric neuroblast division. (**A**) Scheme of a chromosome showing the locations of the *pbl* locus and the Φ31 insertion of the pAcman-pbl genomic constructs indicated on the right. The construct “Pbl-A+B” corresponds to the *pbl* gene and regulatory elements allowing the expression of both Pbl-A and B (Murray et al., 2012a). “Pbl-A” construct corresponds to *pbl* gene and regulatory elements with the non-sense substitution identified in *pbl^MS^*. This allows the expression of Pbl-A but prevents the expression of a functional Pbl-B. “Pbl-B” construct corresponds to *pbl* gene and regulatory elements where exon 8 and 9 are fused to allow the expression of Pbl-B but prevent the expression of Pbl-A. These pAcman insertions were recombined with the *pbl^3^*null mutation. (**B**) Percentage of survival to adulthood of *pbl^3^/pbl^2^*transheterozygote expressing PblA+B, Pbl-A or Pbl-B construct. (**C**) Time-lapse images of neuroblasts of the indicated genotype expressing Sqh::GFP (myosin) from early anaphase to the end of cytokinesis. The images are maximum projections. Time starts at anaphase onset. Time, min:sec. (**D**) Images of neuroblasts of the indicated genotypes expressing Sqh::GFP and the corresponding cortical myosin intensity measurement from apical to basal poles. The black vertical lines represent the position and width of the furrow on the line-scan. (**E**) Scheme illustrating the method to quantify cortical myosin enrichment plotted in (f). (**F**) Scatter dot-plot of cortical myosin enrichment for the indicated genotypes. (**G**) Scheme illustrating the method to calculate the ratio of cortex-to-furrow myosin levels plotted in (h). The blue and red lines correspond to the cortex and furrow respectively. (**H**) Scatter dot plot showing the ratio of cortex-to- furrow myosin levels for the indicated genotypes. (**I**) Selected images of neuroblasts of the indicated genotypes to illustrate the measurement of the area (pink dashed line) and the length (blue line) of the presumptive GMC for the calculation of the roundness plotted in (j). (**J**) Scatter dot plot showing the presumptive GMC roundness for the indicated genotype. (**K**) Graph showing the furrow width over time for the indicated genotypes from anaphase onset to the end of ingression. Data points were fitted to a sigmoid curve (line). Dots and bars indicate mean±SD. n, number of cells. (**L**) Selected DIC images from time-lapse of neuroblasts of the indicated genotypes at the end of cytokinesis. The dashed yellow lines delineate the GMC. (**M**) Scatter dot-plot showing the GMC areas (number of pixels) for the indicated genotypes. Bars represent median ± interquartile range for all scatter dot- plots. A Mann-Whitney two-tailed test was used to calculate P values. Scale bars, 5µm.

To assess the importance of each Pbl isoform on myosin dynamics during neuroblast cytokinesis, we monitored dividing cells expressing the different *pbl* constructs and the myosin marker Sqh::GFP. Live imaging of *pbl^3^/pbl^2^* neuroblasts expressing both Pbl-A and B (Pbl-A+B) revealed that myosin exhibited similar dynamics to wild type cells expressing endogenous Pbl-A and B (presented in Figure 1). Myosin depleted the apical and subsequently the basal pole and accumulated at the baso-equatorial cortex to form the contractile ring. At mid ingression, a pool of myosin invaded the cortex of both nascent cells from the contractile ring to the poles similarly to that of wild type cells (Figure 4C-F, video 3). Importantly, myosin remained more prominently concentrated at the contractile ring than at the polar cortex, as illustrated in myosin intensity profile from pole-to-pole and the ratio of cortex-to-furrow fluorescence intensity for myosin (Figure 4D, G and H). As previously reported, the cortical enrichment of myosin was associated with a slight elongation of the GMC, illustrating an increase in lateral cortical contractility (Figure 4I and J) (Montembault et al., 2017). At the end of furrow ingression, myosin entirely dissociated from the cortex, 3 minutes on average after its initiation of enrichment, as previously reported (Fig. 4c)(Montembault et al., 2017). Importantly, the rate of furrow ingression was similar to wild type cells (compare gray curves in Figure 4K and Figure 1F). Finally, the expression of both isoforms in *pbl* mutant neuroblasts gave similar GMC size as wild type cells (Figure 4L and M, compared gray dots distribution with WT distribution in Figure 1H).

Next we examined myosin dynamics in *pbl^3^/pbl^2^*neuroblasts expressing Pbl-A but lacking a functional Pbl-B. We found that myosin exhibited dynamics reminiscent of those observed in *pbl^MS^* mutant cells. Myosin remained highly focused at the contractile ring throughout furrow ingression (Figure 4C-D, video 3). No cortical enrichment of myosin was detected in any of the cells monitored (Figure 4E-H). Moreover, while the rate of furrow ingression was not affected (Figure 4K), the GMC cell size asymmetry was not preserved. Indeed, the GMCs were significantly smaller in cells expressing only Pbl-A than in cells expressing both isoforms at completion of cytokinesis (Figure 4L and M). These results, first, confirm that the non-sense mutation identified in the *pbl^MS^* sequence is responsible for the perturbed myosin dynamics and the abnormal cell size asymmetry observed in the dividing neuroblasts of the *pbl^MS^*mutant. Second, they indicate that Pbl-B is required for the enrichment of myosin at the polar cortex. Third, they suggest a role for the polar cortex contractility in maintaining ring position during furrow ingression, and more generally in preserving cell size asymmetry.

Next, we monitored myosin dynamics in *pbl^3^/pbl^2^*neuroblasts expressing only Pbl-B and, thus, lacking a functional Pbl-A. Within two minutes after anaphase onset, myosin depleted the poles and enriched the baso-equatorial cortex during contractile ring assembly, as observed for cells expressing Pbl-A only or both isoforms. At mid- furrow ingression, however, myosin displayed a distinct pattern. Myosin spread rapidly toward the polar cortex of both nascent cells (Figure 4C-F, video 3). This was concomitant with its dilution from the contractile ring as illustrated by the lack of maximum levels of myosin at the furrow in the myosin intensity linescan from pole-to-pole and the increase in the ratio of cortex-to-furrow myosin levels (Figure 4D-H). This was accompanied by a severe widening of the furrow, which, in some instances, adopted an extreme tubular shape (Figure 4C, time 3:20 to 5:20, I and J, video 3). In addition, a significant decrease in the rate of furrow ingression was observed (Figure 4K). Despite these altered myosin dynamics and the slower rate of furrow ingression, the GMC cell size was preserved (Figure 4L and M). This wide furrow in Pbl-B expressing cells is reminiscent to the enlarged contractile ring observed when trailing chromatids are present at the cleavage site in wild type cells (Kotadia et al., 2012; Montembault et al., 2017). To test the possibility that the phenotypes observed in Pbl-B expressing cells are due to lagging chromosomes, we examined chromosome segregation concomitantly with myosin dynamics in this genetic background (Fig. S3). No chromosome segregation defect was detected in Pbl-B cells exhibiting a transient widening of the furrow associated with excessive myosin enrichment at the polar cortex (Figure S3A and B).

Changes in myosin dynamics, the rate of furrow ingression and the altered GMC cell size between Pbl-A or B expressing cells could result from the difference in Pbl-A and B localisation, thus inducing distinct patterns of Rho1 activation. Alternatively they could result from a difference in Pbl-A and B function regardless of their localisation. To distinguish between these possibilities, we sought to prevent Pbl-A midzone localisation and examine the outcome in terms of myosin dynamics, furrow ingression and daughter cell size asymmetry. Previous studies have shown that the depletion of RacGAP50C prevents Pbl-A localisation at the furrow during cytokinesis in embryonic cells (Zavortink et al., 2005). Similarly, MgcRacGAP is essential for Ect2 localisation to the midzone of the central spindle in mammals (Yuce et al., 2005). We therefore determined if the Pbl-A midzone localisation in neuroblasts was dependent on RacGAP50C.

Depletion of RacGAP50C by RNAi specifically in imaginal discs and the central nervous system during larval development induced massive death at pupal stage and no survival to adulthood, illustrating the efficiency of the RNAi used to deplete RacGAP50C. In addition, a large number of polyploid cells were observed in third instar larval brains, indicating extensive cytokinesis failure in this tissue (Figure S4) as previously reported (Roth et al., 2015). However, in the course of our live study described below, only 2 out of 38 RacGAP50C depleted neuroblasts failed cytokinesis during the recording. One explanation for this discrepancy between the frequencies of polyploidy in fixed brains and cytokinesis failure during live imaging is that most of the cytokinesis defects in RacGAP50C depleted cells may occur at later stages after termination of furrow ingression, and, hence, after our live recording. Consistently, depletion of centralspindlin components in Drosophila neuroblasts or Cyk-4 in the *C. elegans* one-cell embryo does not affect furrow ingression but results in cytokinesis failure at a later stage (Jantsch-Plunger et al., 2000; Roth et al., 2015).

Next, we monitored GFP::Pbl-A dynamics in neuroblasts depleted for RacGAP50C by RNAi. We found that, while GFP::Pbl-A signal was not detected at the midzone and was not enriched at the site of furrowing, its broad localisation at the nascent cells plasma membrane was not affected (Figure 5A). We conclude that RacGAP50C is required for Pbl-A midzone and furrow localisation but is not essential for its recruitment to the plasma membrane. Remarkably, these RacGAP50C depleted cells exhibited transient enlarged furrows reminiscent of those observed in cells expressing Pbl-B.

**Figure 5.**
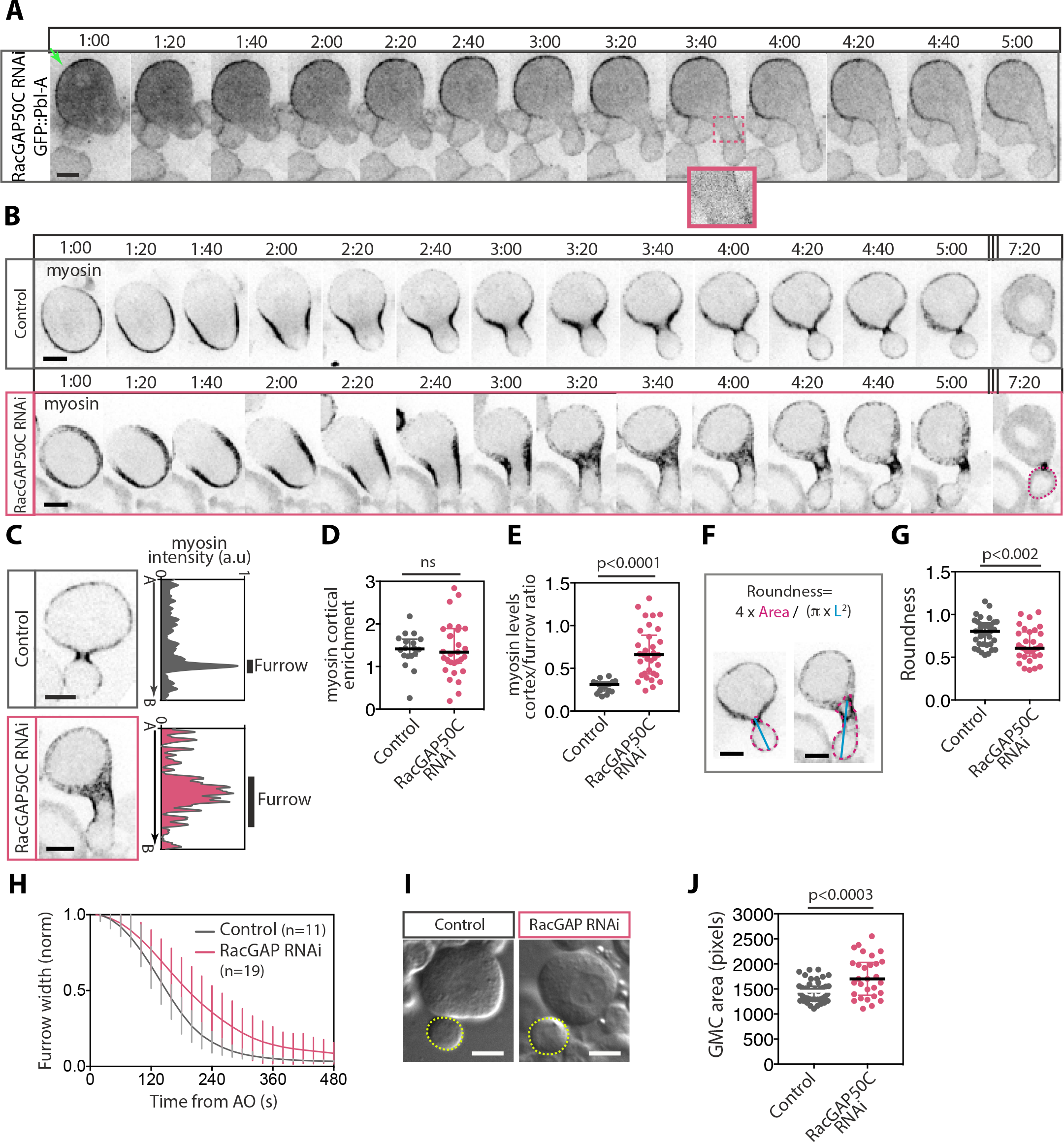
Cytokinesis defects in RacGAP50C depleted neuroblasts are similar to that observed in cells lacking Pbl-A. (**A**) Time-lapse images of neuroblasts depleted of RacGAP50C by RNAi and expressing GFP::Pbl-A. Inset shows the magnified midzone and furrow region delineated by the dashed pink square on the image. GFP::Pbl-A is readily detected on the nascent daughter cells plasma membrane but not at the midzone in RacGAP50C depleted cells. (**B**) Time-lapse images of control neuroblasts or neuroblasts depleted of RacGAP50C by RNAi (RacGAP50C RNAi) expressing Sqh::GFP (myosin) from early anaphase (one minute after anaphase onset) until the end of furrow ingression. RacGAP50C depleted cell exhibited an enlarged furrow associated with a dilution of myosin from the furrow and an excessive enrichment of myosin on the polar cortex. Time starts at anaphase onset. (**C**) Selected images of neuroblasts of the indicated genotypes expressing Sqh::GFP (myosin) and the corresponding cortical myosin intensity measurement at the cortex from apical-to-basal poles. The vertical lines indicate the position and width of the furrow on the line-scan. (**D**) Scatter do-plot showing cortical myosin enrichment for the indicated genotypes. (**E**) Scatter dot-plot showing the ratio of cortex-to-furrow myosin levels of the indicated genotypes (as described in Figure 4G). (**F**) Selected images of neuroblasts to illustrate the area (pink dashed line) and the length (blue line) measured for the calculation of the roundness of the presumptive GMC plotted in (g). (**G**) Scatter dot-plot showing the roundness of the presumptive GMC for the indicated genotype at the time of maximum deformation. (**H**) Graph showing the furrow width over time for the indicated genotypes from anaphase onset to the end of ingression. Data points were fit to a sigmoid curve (line). Bars indicate mean±SD. n, number of cells. (**I**) Selected DIC images from time-lapse of neuroblasts of the indicated genotypes at the end of cytokinesis. The dashed yellow lines delineate the GMC. (**J**) Scatter dot-plot showing the GMC area (number of pixels) for the indicated genotypes. Bars represent median ± interquartile range for all scatter dot-plots. A Mann-Whitney test was used to calculate P values. Scale bars, 5µm. Time, min:sec.

This prompted us to examine myosin dynamics, the rate of furrow ingression and GMC cell size upon RacGAP50C depletion. Since depletion of RacGAP50C results in a dramatic increase in the number of polyploid neuroblasts (Figure S4), we chose to examine the division of diploid neuroblasts only, to exclude phenotypes that could result from polyploidy. These diploid neuroblasts can be readily distinguished from their polyploid counterparts by their diameter at metaphase and the size of the chromosome mass. Control cells exhibited myosin dynamics similar to that reported for wild type cells throughout cytokinesis (Figure 5B-E, video 4). In contrast, attenuation of RacGAP50C altered myosin dynamics in a similar way as cells expressing only Pbl-B. At mid-ingression, myosin formed an unusually large ring that produced an abnormally wide furrow (Figure 5B, C, F and G, video 4). Subsequently, myosin underwent excessive cortical enrichment as illustrated by the high ratio of cortex-to-furrow myosin levels compared to control cells (Figure 5C-G). The broadening of myosin cortical localisation was concomitant with a decrease in the rate of furrow ingression (Figure 5H). These cytokinesis defects were associated with a mild proportion of larger GMCs than normal at the end of furrow ingression (Figure 5J). Since RacGAP50C depleted cells share similar phenotypes as Pbl-B expressing cells, including excessive myosin cortical relocalisation, enlarged furrow and a slower rate of furrow ingression, we conclude that these phenotypes are not the result of the absence of Pbl-A but rather the lack of midzone and furrow-localized Pbl-A signalling which favours Rho1 activation through broad Pbl-B membrane localisation.

## Discussion

In this study we provide evidence that the asymmetric division of the drosophila neural stem cell relies on the activity of two Pbl isoforms displaying distinct localisation patterns. These two isoforms modulate Rho1 activity concurrently to promote robust furrow ingression while allowing the adjustment of its asymmetric position during ring closure. Based on our findings, we propose the following model: Pbl-A, predominantly concentrated at the midzone and furrow site through its association with RacGAP50C, maintains a threshold of Rho1 signalling at the leading edge of the furrow to sustain efficient ingression through myosin activation. After initiation of furrowing, the broad localization of Pbl-B at the plasma membrane enlarges the zone of Rho1 activity, which, in turn, drives myosin efflux from the contractile ring and its enrichment at the polar cortex. This increase in poles contractility contributes to the fine-tuning of furrow position to preserve daughter cell size asymmetry. When Pbl-B is absent, Pbl-A restricts Rho1 activity in a narrow zone, which is sufficient to drive efficient furrow ingression but is insufficient to adjust furrow position. Consequently, abnormally small GMCs are produced, depending on the initial position of the furrow. When Pbl-A is not enriched at the midzone and furrow, Pbl-B-dependent broadening of Rho1 activity dilutes active myosin from the edge of the furrow by promoting its excessive enrichment at the polar cortex. As a consequence, the furrow enlarges which temporarily diminishes its rate of ingression (Figure 6).

**Figure 6.**
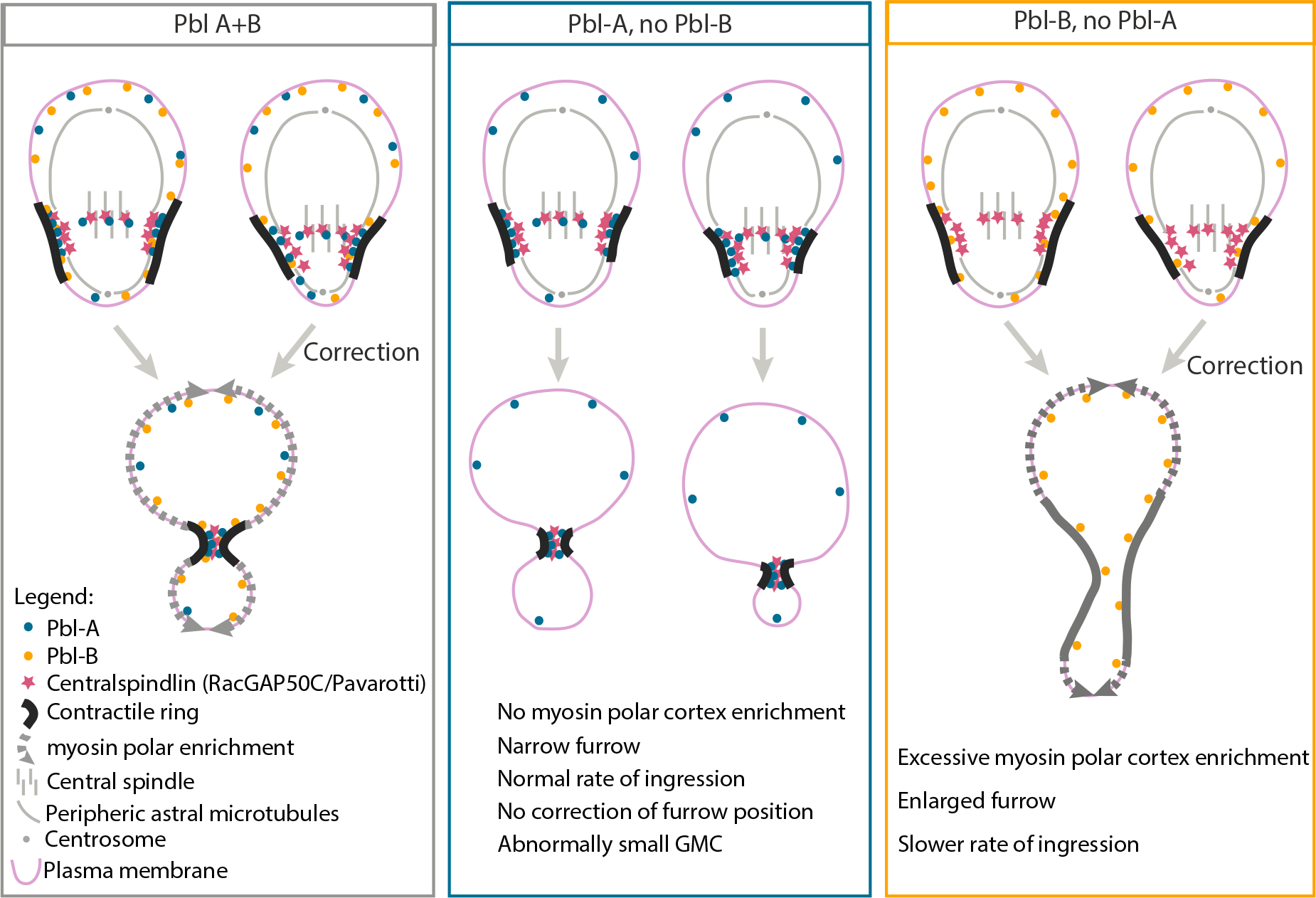
Model for Pbl-A and B function during neuroblast asymmetric division. Two Pbl isoforms, Pbl-A and B with distinct localisation patterns act concomitantly to promote robust asymmetric cytokinesis. Pbl-A (blue circle) remains predominantly at the midzone and site of furrowing through its association with the centralspindlin component RacGAP50C (pink star), which interacts with two pools of microtubules, the peripheral astral microtubules and the central spindle. The enrichment of Pbl-A at the furrow sustains a threshold of Rho1 activity to promote efficient ring constriction. Pbl-B localizes broadly on the plasma membrane where it enlarges the zone of Rho1 activity throughout the second phase of furrow ingression. This promotes polar cortical myosin enrichment on both nascent daughter cells. The balance between the activities of these two Pbl-isoforms allows efficient furrow ingression while adjusting contractile ring position to generate correct unequal-sized daughter cells. In cells expressing Pbl-A but not Pbl-B, Rho1 activity remains focused at the furrow throughout ingression. As a result, myosin does not enrich the polar cortex. This focused Rho1 activity permits efficient ring constriction but prevents adjustment of furrow position to preserve the asymmetry in cell size. Consequently, abnormally small GMCs are produced in the instance where the furrow is initially too basally positioned. In cells expressing Pbl-B but not A, Pbl-B broad membrane localisation promotes Rho1 activation in a larger zone after furrow initiation. Consequently, myosin undergoes excessive polar cortex enrichment. This induces the transient widening of the furrow accompanied by a decrease in its rate of ingression. Normal sized GMC are nevertheless produced at the end of furrowing.

### Pbl-B is one of the two major Pbl isoforms with functions during cytokinesis and development

The identification of the *pbl^MS^* mutation as a non-sense substitution in the alternatively spliced exon 8 led us to uncover Pbl-B as one of the two major Pbl protein isoforms expressed in all drosophila tissues. The premature stop codon in exon 8 prevents the expression of a functional Pbl-B in the *pbl^MS^* mutant.

No role has been assigned to Pbl-B isoform prior to our work. Two distinct functions for Pbl during embryonic development have been identified through genetic screens: (1) its function as a GEF for Rho1 during cytokinesis (Hime and Saint, 1992; Lehner, 1992; Prokopenko et al., 1999; Somers and Saint, 2003) and (2) its Rho1- independent role in the epithelial to mesenchymal transition (EMT) and mesoderm cell migration (Murray et al., 2012a; Murray et al., 2012b; Nakamura et al., 2013; Schumacher et al., 2004; Smallhorn et al., 2004; van Impel et al., 2009). The fact that the expression of Pbl-B rescues the embryonic lethality of *pbl* null mutants and produces adult escapers indicates that it contributes to some extent redundantly with Pbl-A to perform these two independent functions.

### An alternative, RacGAP50C-independent pathway, for Pbl activation during cytokinesis

Structural modelling predicts that Pbl-B, like Pbl-A, may possess three folded BRCTs on its N terminus despite the presence of 457 extra amino acids within the third BRCT sequence. Since BRCTs are critical for Pbl-A interaction with RacGAP50C, this implies that Pbl-B may bind RacGAP50C. However, several lines of evidence suggest that, *in vivo*, Pbl-B functions independently of its interaction with RacGAP50C, unlike Pbl-A. First, Pbl-B is not detected at the spindle midzone and is not concentrated at the furrow, whereas Pbl-A localizes to the midzone and furrow site in a RacGAP50C-dependent manner. Second, expression of only the Pbl-B isoform produces phenotypes, including an enlarged furrow and slower rate of ingression, akin to that observed in cells depleted for RacGAP50C. Hence, we propose that two parallel pathways, RacGAP50C-dependent Pbl-A and RacGAP50C- independent Pbl-B pathways act in concert to promote robust neuroblast asymmetric division. Consistently, studies in *C. elegans* one-cell embryos revealed that two parallel pathways, a Cyk-4-dependent and independent pathways, drive furrow ingression (Jantsch-Plunger et al., 2000; Tse et al., 2012). The Cyk-4-independent pathway utilizes the non-conserved NOP1 protein for Ect2 activation (Tse et al., 2012). In human cultured cells, the association of Ect2 with the plasma membrane or the local activation of RhoA through the use of optogenetics is sufficient to induce furrowing (Kotynkova et al., 2016; Wagner and Glotzer, 2016). Remarkably, successful cytokinesis occurs in cells expressing Ect2 mutated on residues that impair its interaction with MgcRacGAP and prevent its recruitment to the spindle midzone, highlighting an MgcRacGAP-independent redundant pathway for Ect2 activation (Kotynkova et al., 2016). Recently, a novel isoform of human Ect2 has been identified through a screen for mammalian transformed cells resistant to treatment with Doxorubicin (Tanaka et al., 2020). Importantly, this long human Ect2 isoform is strictly nuclear in interphase like Pbl-B and, like Pbl-B, has additional amino acids in one of the BRCT domains that might affect its proper folding and impair Ect2 interaction with MgcRacGAP. Thus, the use of redundant RacGAP- dependent and independent pathways for Ect2/Pbl activation during cytokinesis may be a convergent mechanism in multicellular organisms, providing robustness to a process that has to adapt to a variety of cell types with specific extrinsic and intrinsic physical constraints.

The model that Pbl-B drives cytokinesis independently of RacGAP50C raises the question of how the furrow site is specified in these cells. Previous studies have identified two distinct pathways that define the future cleavage site in Drosophila neuroblasts. One pathway involving the Pins polarity determinant complex acts early to promote myosin depletion from the apical pole and accumulation at the basal cortex concomitant with the expansion of the apical membrane (Cabernard et al., 2010; Connell et al., 2011; Pham et al., 2019; Roubinet et al., 2017). The second pathway depends on microtubules-associated centralspindlin complex that acts to stabilize the baso-equatorial position of the contractile ring to the presumptive cleavage site (Roth et al., 2015; Thomas et al., 2021). Since myosin dynamics during furrow site specification are unperturbed in cells lacking Pbl-A or depleted for RacGAP50C, we propose that the polarity-determinant pathway may be the predominant pathway that positions the cleavage site in these cells. The target of the polarity pathway for cleavage site specification is currently unknown. It would be interesting in the future to determine if this pathway acts through Pbl-B regulation.

### Mechanics of cell size asymmetry

One of our most striking findings was that cells without Pbl-B produce aberrantly small GMCs. The fact that the GMC size is correlated with the initial position of the furrow in these cells suggests a default in furrow position correction. Previous studies have determined that two pool of centralspindlin complex spatially separated by two different populations of microtubules (the central spindle and the peripheral astral microtubules), competitively influence furrow position during ingression (Thomas et al., 2021). In this study they showed that perturbation of peripheral microtubules-associated pool of centralspindlin produces larger GMCs than normal. Since our data suggest that Pb-B acts independently of the centralspindlin complex and that cells lacking Pbl-B produce GMC that are smaller than normal, we propose that an additional mechanism than the centralspindlin-dependent pathway participates in furrow adjustment. We can formulate hypotheses based on our findings and those of others. The accumulation of myosin at the poles during furrow ingression is likely to change the mechanical properties of the nascent daughter cell cortices (Reichl et al., 2005). Indeed, myosin enrichment at the polar cortex coincides with an increase in contractility, as illustrated by the transient deformation of the nascent cells (Fig. 4j). Thus, we speculate that the increase in contractility of the neuroblast cortex following myosin enrichment may produce sufficient compressive force to drive cytoplasmic advection towards the GMC, thereby adjusting furrow position by inducing GMC swelling. This implies an anisotropic localisation of active myosin in favour of the large nascent cell cortex. A prediction of this model is if a compressive force could be applied on the nascent neuroblast by micromanipulation, this should rescue the small GMC phenotype in the Pbl-B depleted cell. Support for this model comes from studies in human cultured cells demonstrating that perturbation of the contractile properties of one nascent daughter cell polar cortex induces furrow displacement associated with cytoplasmic advection and cell shape instability (Sedzinski et al., 2011; Yamamoto et al., 2021). Furthermore, the polarized myosin activity on one daughter cell cortex is critical to produce unequal-sized daughter cells during the division of *C. elegans* Q lineage neuroblasts (Ou et al., 2010). Finally, studies in mammalian cells have shown that two myosin isoforms with different motor kinetics participate in polar cortex contractility, which is critical for maintaining cell shape stability and promoting the fidelity of cytokinesis (Taneja et al., 2020; Yamamoto et al., 2019). Alternatively, the myosin-deprived polar cortex of Pbl-B depleted cells may be more elastic than myosin-enriched WT polar cortex, and, thus, may favour cytoplasmic advection from the GMC to the neuroblast due to the increasing difference in hydraulic pressure between the small and large nascent daughter cells as the furrow ingresses. This will be exacerbated by the extent to which the initial site of furrowing is positioned on the basal side.

Our work provides ground for future studies on how changes in the mechanical properties of the polar cortex contribute to adjust furrow position during ingression. Pbl-A and B isoforms provide excellent genetic tools to address this question as they allow the physiological modulation of Rho1 activity at the furrow and polar membrane. It will be equally interesting to assess this question in relation to tissue features.

## Acknowledgments

We thank Andrew Murray for sharing reagents. We thank Régis Giet, Gilles Hickson and Cameron Mackereth for helpful discussion. We thank Marie Didelon for technical help and all lab members for insightful discussion. EM, LB, DMcC and AR were supported by Centre National de la Recherche Scientifique (CNRS). AR was also supported by the European Research Council (ERC-GA311358), the Agence Nationale de la Recherche (ANR-n°234520), and the Conseil Régional d’Aquitaine (2014-1R30412-00003094). MCC was supported by the University of Bordeaux. ID was supported by the Fondation pour la Recherche Médicale (FRM- ECO202106013724) and CNRS.

## Methods

### Fly stocks

Flies were raised on standard media at 25°C. The *pbl^2^*and *pbl^3^* alleles were described previously (Jurgens et al., 1984; Prokopenko et al., 1999; Somers and Saint, 2003). The screen that identified *pbl^MS^* (also called pblZ4836) was described in (Giansanti et al., 2004). P[UAS>RacGAP50C-dsRNA] (stock n°6439) and P[69B>Gal4] strains were provided by Bloomington stock center (indiana, USA). P[w+, sqh>Sqh::GFP transgenic strain was described in (Royou et al., 2004; Royou et al., 2002). P[UAS>VenusFP::RacGAP50C] was described in (Goldstein et al., 2005). Pbl-A+B stock was described in (Murray et al., 2012a). Pbl-A, Pbl-B, GFP::Pbl-A and GFP::Pbl-B transgenic stock were produced in our laboratory for this study (see Table S1 for the exact genotype of flies used in this study).

### Sequencing of pebble genomic region

Genomic DNA was extracted from 30 adult flies previously collected and frozen at - 80°C, by grinding in 400µL of buffer containing 100mM Tris-HCl pH7.5, 100mM EDTA, 100mM NaCl, 0,5% SDS, until only cuticles remain. The mixture was incubated for 30 minutes at 65°C and 800µL of a solution of LiCl/KAc (1 part of 5M KAc:2.5 parts 6M LiCl) was incubated on ice for 10 min. After centrifugation at 12000g for 15 minutes, 1mL of supernatant was transferred in 600µL of isopropanol. After 15 minutes of centrifugation at 12000g, the pellet containing the genomic DNA was washed with 70% ethanol and dried before resuspension in 150µL of diH2O.

Amplification by PCR of three fragments covering the whole genomic region of pebble was performed using oligonucleotide pairs A to C from genomic DNA of WT and homozygous *pbl^MS^*flies (Table S2). Each fragment was sequenced for complete coverage of exons in the pebble gene. Sequences were analyzed with Serial Cloner software and compared to Flybase annotated genome (FBgn0003041).

### RT-qPCR

To estimate the level of pebble mRNA isoforms in larval brains, total RNAs from 60 larval brains were extracted using RNAspin Mini kit (GE Heathcare). 1µg of total RNAs were converted into cDNA using oligodT and SuperScript III Reverse transcriptase (Invitrogen) according to the manufacturer’s protocol. Quantitative PCR was performed using SSO Advanced Universal SYBR Green Supermix (Bio-Rad).

### Cloning

Pbl-A and B constructs were obtained by DNA synthesis using pAcman[Pbl] plasmid described previously (Murray et al., 2012a). For Pbl-A, the cytosine at position 1114 (Q372) was substituted to a thymidine to mimic the *pbl^MS^*mutation. For Pbl-B, the intron sequence between exons 8 and 9 was removed (Genebridges). GFP::Pbl-A and GFP::Pbl-B were cloned as followed: Pbl-A and B cDNAs produced by RT-PCR from adult mRNA extracts were inserted into pAcman[GFP] using AscI and NotI sites flanking a cassette obtained by PCR and containing sequences as followed: AscI-pbl promoter-pbl5’UTR-GFP-pbl (A or B) cDNA -pbl3’UTR-NotI. The resulting plasmids pAcman[Pbl-A], pAcman[Pbl-B], pAcman[GFP::Pbl-A] and pAcman[GFP::Pbl-B] were used to produce transgenic flies with the insertion at the same landing site, 86F8b on the third chromosome. The plasmid pAcman[GFP::Pbl-A+B] was produced by inserting the eGFP sequence before the ATG of the genomic Pbl sequence present in pAcman[Pbl] using AscI and BsiW restriction enzymes. The resulting pAcman[GFP::Pbl] plasmid was inserted at the landing site 68A4 on the third chromosome of the fly genome.

### Western-blot

60 brains from 3^rd^ instar larvae of each genotype were dissected in PBS and transferred in 30µL of RIPA buffer (Tris-HCl 10mM pH7.5, NaCl 150mM, EDTA 0.5mM, SDS 0.1%, Triton 1%, Na Deoxycholate 1%) supplemented with protease inhibitor cocktail (Roche) and PMSF (1mM). After grinding and homogenization with a pestle in a 1.5mL centrifuge tube, lysates were centrifuged at 15000g for 5min at 4°C and supernatants containing proteins were transferred into equal volume of Laemmli 2X buffer. Boiled samples were loaded and analyzed by 10% SDS-PAGE. Proteins were transferred to nitocellulose membrane using a semi-dry electrophoretic blotting device (BioRad). Mouse anti-GFP (Roche, clones 7.1 and 13.1, 2µg/ml) and mouse anti-αTubulin primary antibodies (Sigma, clone DM1A, 2µg/ml) and anti-mouse HRP conjugated secondary antibodies (DakoCytomation, 2µg/ml) were used for western blotting.

### Microscopy

Live imaging of larval neuroblasts was carried out as previously described (Landmann et al., 2020). Late third instar larval brains were dissected in PBS and transferred to a drop of PBS on a coverslip. The brain was slightly squashed between a slide and coverslip by capillary force to facilitate visualization of dissociated neuroblasts. The coverslip was then sealed with halocarbon oil 700 (Sigma). In this study, only neuroblasts between 11 and 12 µm in diameter were observed. Live analysis was performed at room temperature with 100X oil Plan-Apochromat objective lens (NA 1.4) and an Axio-Observer.Z1 microscope (Carl Zeiss) equipped with a spinning disk confocal CSU-X1 (Yokogawa), an EMCCD Evolve camera (Photometrics). The fluorescent proteins were excited with a 491nm (100mW; Cobolt Calypso) and 561nm (100mW; Cobolt Jive) lasers. Metamorph software (GATACA) was used to acquire the data. Images were acquired every 20 seconds over 7µm depth with a z step of 0.5µm. Images are maximum projections of selected z unless specified in the figure legend.

### Image analysis and data processing

Fiji (National Institute of Health) was used for image analysis and fluorescence quantification (Schindelin et al., 2012). The background level of the camera was substracted from all raw data before quantification. For the line-scan in Figures 1C, 4D and 5C, a 2-pixel wide line was drawn around the cortex from one cell pole to the other at the time of maximum myosin polar cortical enrichment (∼80% furrow ingression). The average gray value in the cytoplasm was substracted from the data. The maximum gray value was used for normalization and the line-scan was plotted (6 neighbors average) using prism (GraphPad). For Figures 1E, 4F and 5D, the average gray value of myosin at the neuroblast cortex (Myo_cortex_) and the cytoplasm (Myo_cyto_) was measured using a 2-pixel wide line as illustrated in the schemes (Figure 1D and 4E). Myosin cortical enrichment was subsequently calculated as follows, (Myo_cortex_- Myo_cyto_)/ Myo_cyto_. For Figures 4H and 5E, the mean gray value of myosin at the furrow (Myo_furrow_) was measured by drawing a 2-pixel wide line at the furrow. The length of the furrow was determined by the divergent angle from the long axis of the cell being approximately <30°. The ratio Myo_cortex_/Myo_furrow_ was then calculated. For Figure 1I, the time of initiation of furrow ingression was determined as the time of curvature. The position of the furrow was defined as the maximum point of curvature.

### Viability analysis

For viability assays, flies were raised on standard media at 25°C. Eight young males *pbl^2^*/TM6B were crossed with ten young females of the following genotypes: *pbl^3^*, pAcman[Pbl-A+B]/TM6B, *pbl^3^*, pAcman[Pbl-A]/TM6B, *pbl^3^*, pAcman[Pbl-B]/TM6B or *pbl^3^*/TM6B. For each genotype, the parents were left to lay eggs in a vial every 2 days for 8 days. The first vial was discarded and all the adult progeny of the remaining vials were counted. The percentage of viability of the transheterozygous individuals carrying the pAcman Pbl constructs was determined by calculating the ratio of *pbl* transheterozygote vs TM6B flies over the expected frequency of transheterozygotes according to Mendel’s law of inheritance. The experiment was performed three times independently.

### Rapid Iterative Negative Geotaxis (RING) assay

The Rapid Iterative Negative Geotaxis (RING) assay described by Gargano et al. 2005 (Gargano et al., 2005) was modified as follows: young wild-type and *pbl^MS^* males were collected under CO_2_ anesthesia and allowed to recover in a fresh vial containing food for 5 hours at 25°C. The flies were then transferred without anesthesia into new vials without media and left to acclimatize for 15 minutes at room temperature. The vials were then placed into the ring apparatus, composed of a polystyrene rack with calibrated graduation behind the vials. The vials in the rack were manually tapped onto the surface of a bench three times. This cycle was repeated 5 times every 30 seconds. The experiment was recorded by video and the average height climbed 3 seconds after the last tap was scored for each cycle. The experiment was performed five times independently.

### Longevity test

For each genotype, 5 males and 5 females were collected at birth and placed in a vial with standard media and raised at 25°C. The adults were flipped into fresh media every four days without the use of CO2. The vials were scored for dead flies every day. The experiment was repeated 5 times.

## Supplemental informations

### Supplemental method

#### Cytology

Third instar larval brain were dissected in PBS and stained with aceto-orcein as in (Karess and Glover, 1989). To quantify diploid and polyploid mitotic cells, preparations were observed by phase contrast using a Nikon eclipse Ti equipped with 100X objective.

## Supplemental figures

**Figure S1.**
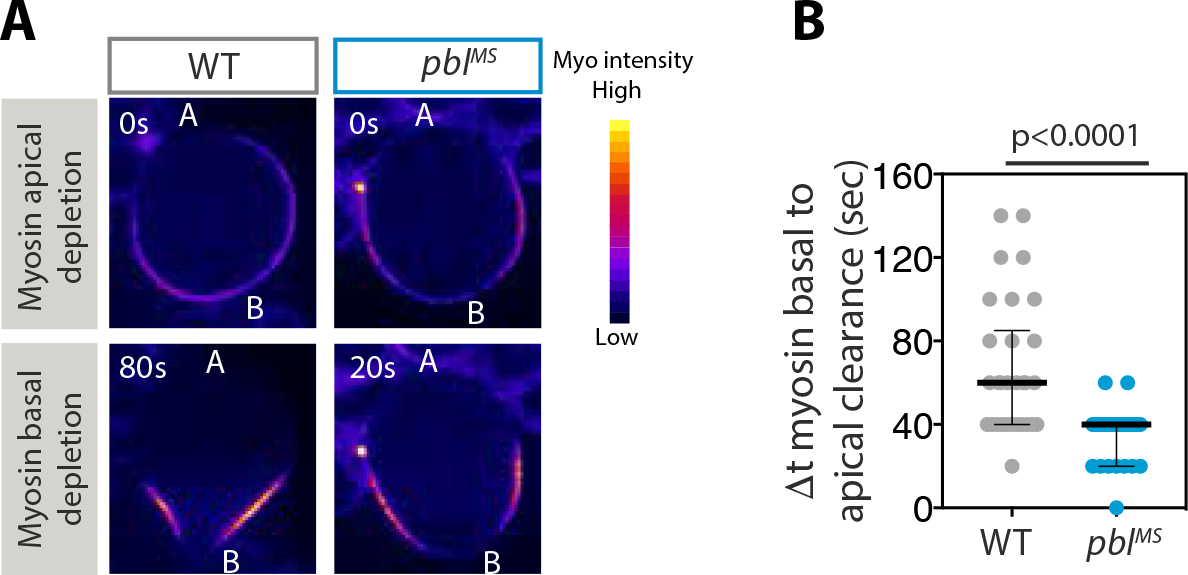
The time elapsed between myosin apical and basal depletion is longer in wild-type than *pbl^MS^*neuroblasts. (**A**) Images of wild-type and *pbl^MS^* neuroblasts expressing Sqh::GFP (myosin) at onset of myosin depletion from the apical (A) and basal (B) poles (top and bottom rows respectively). Time, seconds (s). Scale Bar, 5µm. (**B**) Scatter dot plot showing the time elapsed between myosin apical and basal pole depletion for the indicated genotype. Bars represent median ± interquartile range. A Mann-Whitney test was used to calculate the P value.

**Figure S2.**
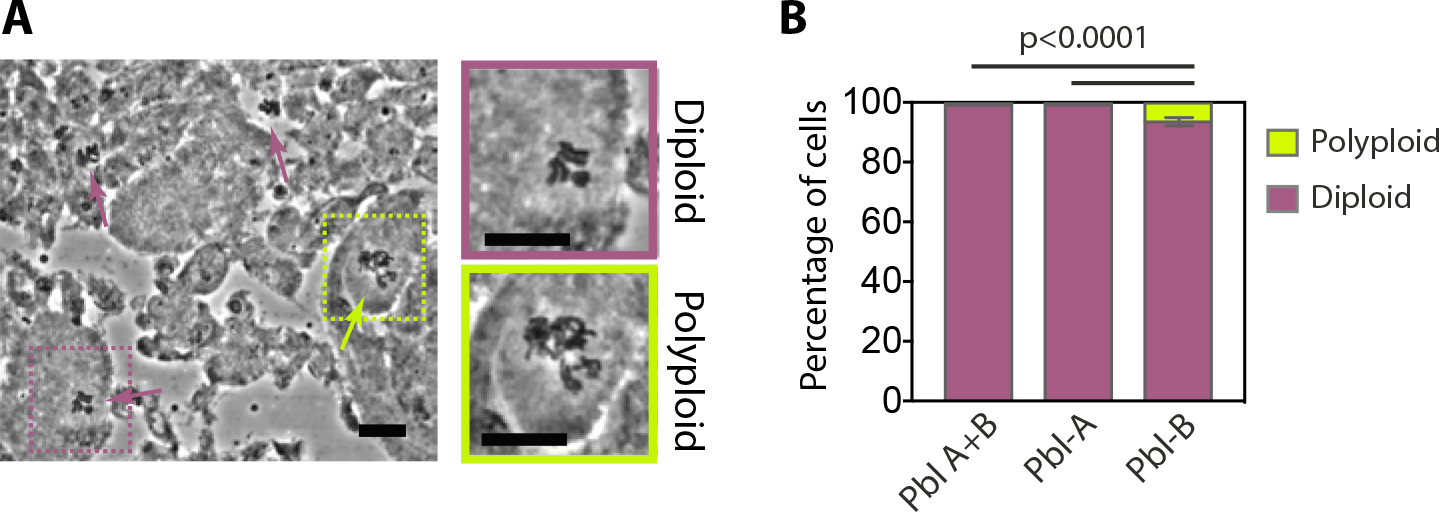
Mild increase in the frequency of polyploid cells in the central nervous systems of third instar larvae expressing solely the Pbl-B isoform. (**A**) Phase contrast image of a squashed larval central nervous system stained with orcein. The purple and yellow arrows point to diploid and polyploidy cells respectively. The insets are magnification of diploid and polyploidy cells. Scale bar, 10µm. (**B**) Frequency of polyploid mitotic cells for the indicated genotype. More than 140 cells were counted per brain and 3 to 4 brains were counted per experiment. Three independent experiments were performed. The bars represent the mean±SD.

**Figure S3.**
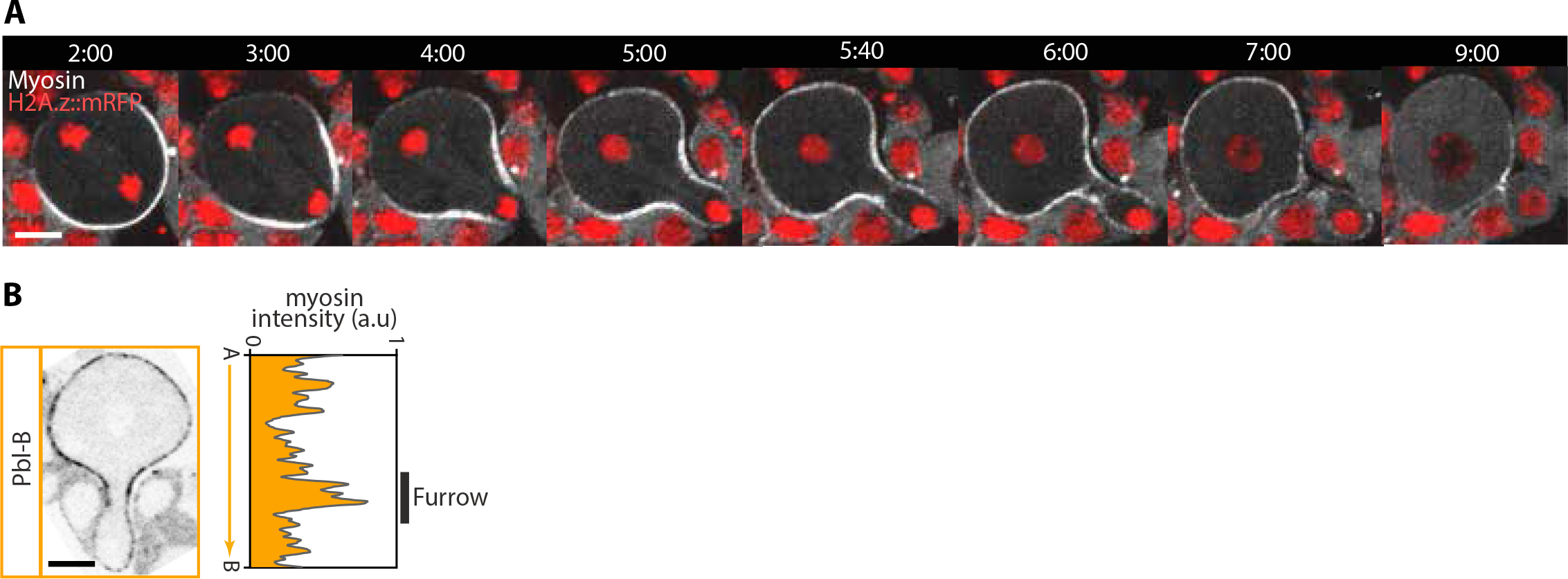
No trailing chromatids are observed in Pbl-B expressing cells. Time-lapse images of Pbl-B expressing cells labelled with H2A.z::mRFP and Sqh::GFP (Myosin). Time starts at anaphase onset. Time, min:sec. Scale bar, 5µm.

**Figure S4.**
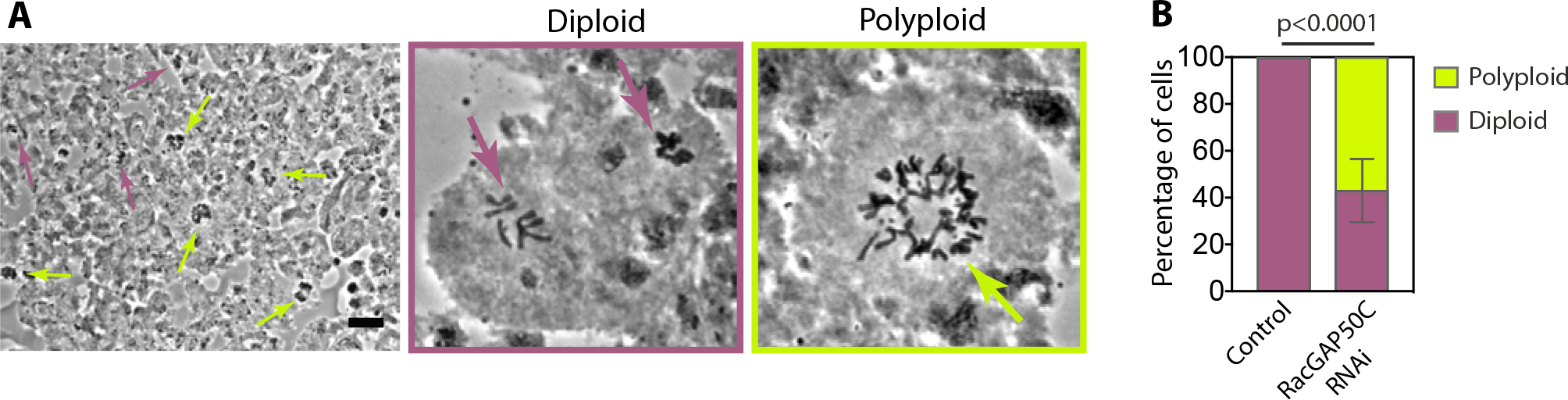
Dramatic increase in polyploid cells upon RacGAP50C depletion in the third instar larvae central nervous system. (**A**) Phase contrast image of a squashed larval central nervous system stained with orcein. The purple and orange arrows indicate diploid and polyploidy cells respectively. The insets are magnification of diploid and polyploidy cells. Scale bar, 10µm. (**B**) Frequency of polyploid mitotic cells for the indicated genotype. More than 80 cells were counted per brain. 4 and 10 brains were counted for control (69B>Gal4) and RacGAP50C RNAi (69B>Gal4; p[RacGAP50C dsRNA]) respectively, in two independent experiments. The bars represent the mean±SD.

## Supplemental videos

Video 1. Myosin dynamics during cytokinesis in wild-type and *pbl^MS^* **neuroblasts.**

Time-lapse video of wild-type (left) and *pbl^MS^* (right) neuroblasts expressing Sqh::GFP. Images are maximum projections. Time=min:sec. Time 0:00 corresponds to anaphase onset (initiation of sister chromatid separation). The movie corresponds to Figure 1A.

Video 2. RacGAP50C, Pbl-A and Pbl-B dynamics during cytokinesis.

Time-lapse video of neuroblasts expressing VenusFP::RacGAP50C (Left), GFP::Pbl- A (middle) and GFP::Pbl-B (Right) from anaphase onset. The images are maximum projections. Time=min:sec. The movie corresponds to Figure 3.

**Video 3. Myosin dynamics during cytokinesis in Pbl-A+B, Pbl-A or Pbl-B expressing cells.** Time-lapse video of *pbl^3^*/*pbl^2^*neuroblasts expressing Pbl-A and B, Pbl-A or Pbl-B and Sqh::GFP. Images are maximum projections. Time, 0:00 corresponds to anaphase onset. Time, min:sec. The movie corresponds to Figure 4C.

**Video 4. Myosin dynamics during cytokinesis in control neuroblasts and neuroblasts depleted for RacGAP50C.** Time-lapse video of 69B>Gal4 neuroblasts (left) and RacGAP50C RNAi; 69B>Gal4 (Right) neuroblasts expressing Sqh::GFP. Images are maximum projections. Time, 0:00 corresponds to anaphase onset. Time, min:sec. The movie corresponds to Figure 5B.

## Supplementary Tables

**Table S1.**
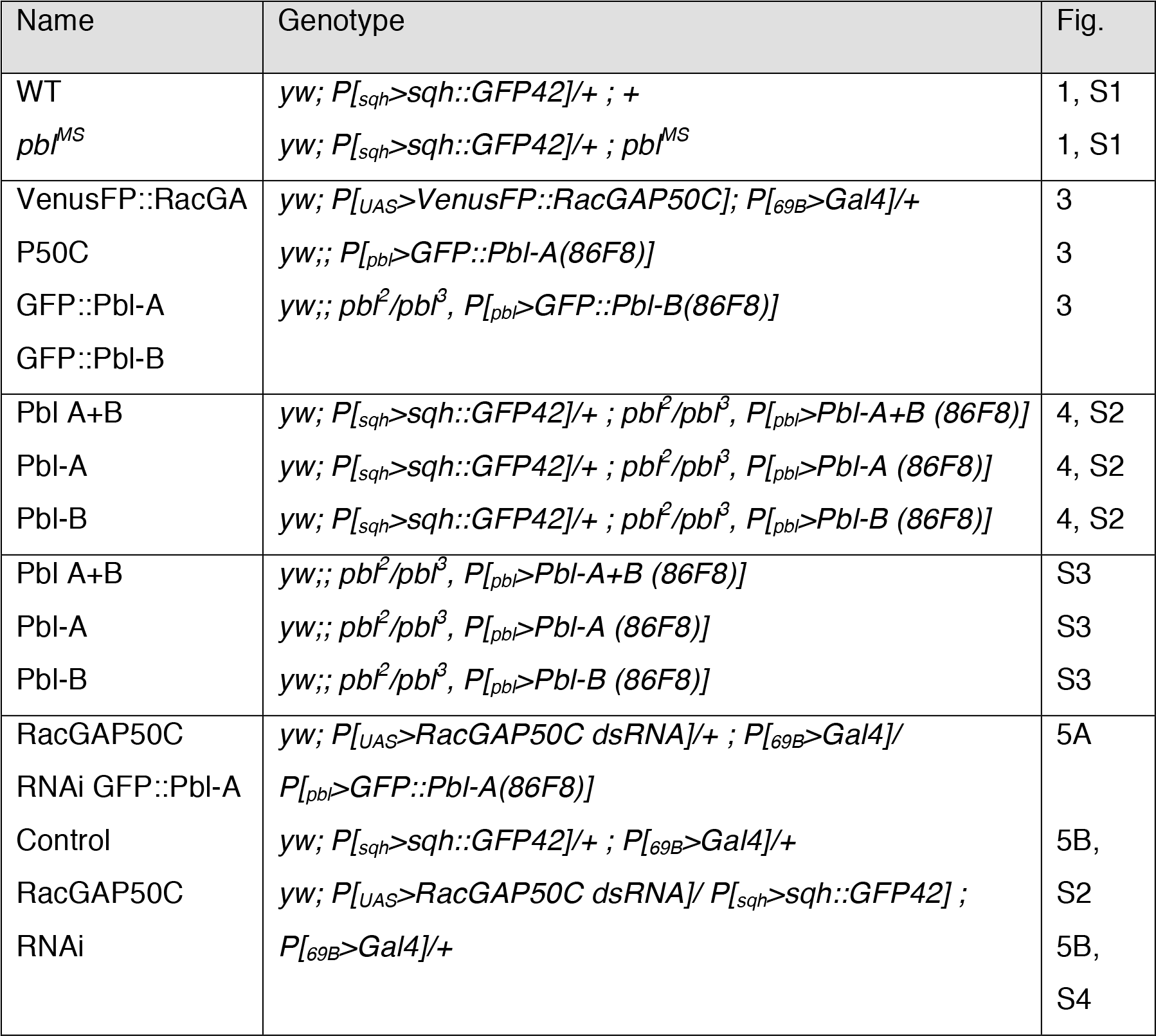

**Table S2.**
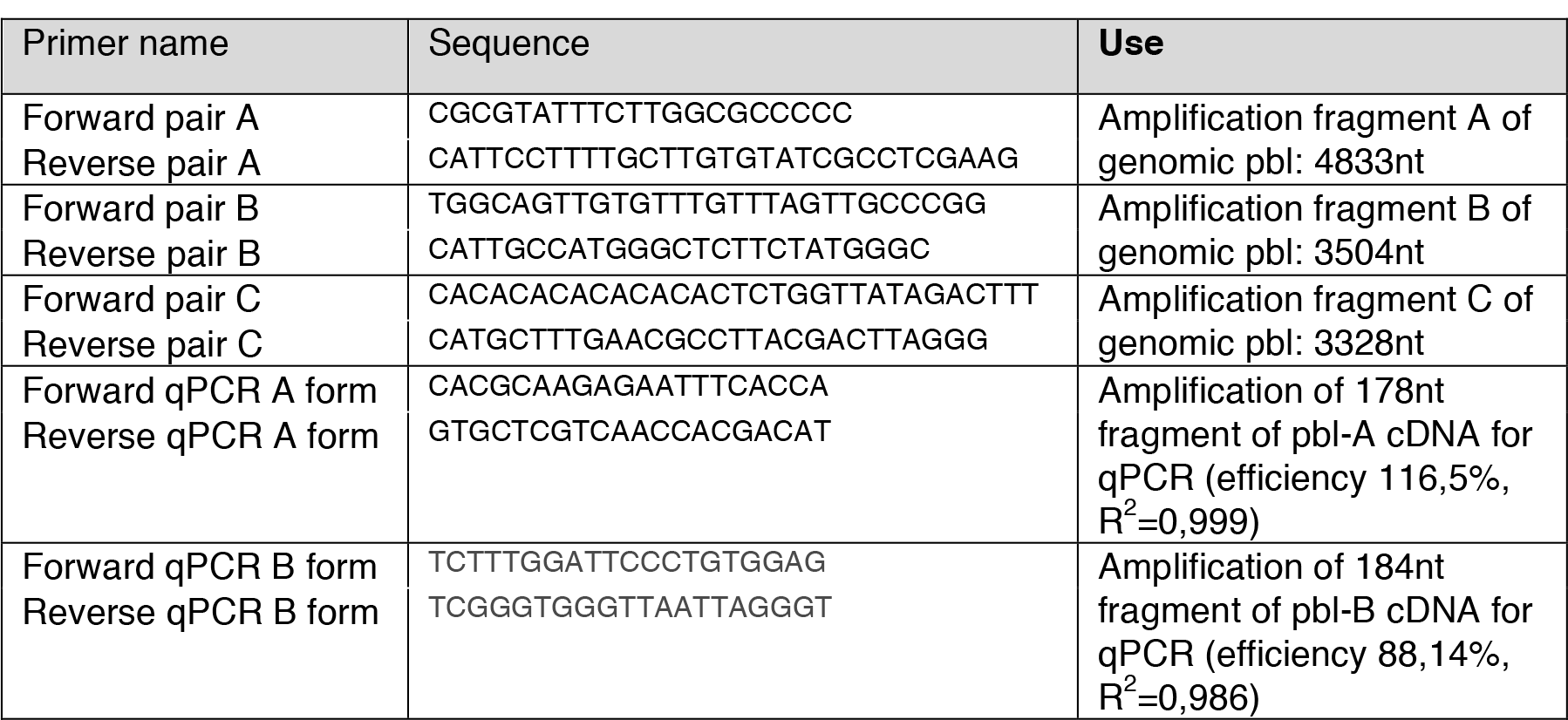
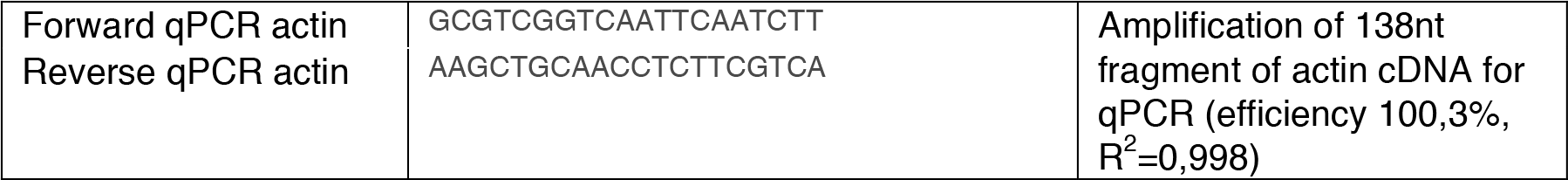

